# When knowledge interferes with perception: Neural mechanisms of the semantic amplification of visual false memory

**DOI:** 10.64898/2026.02.24.707651

**Authors:** Loris Naspi, Şafak Erener, Simon W. Davis, Roberto Cabeza

## Abstract

Visual false memory refers to our tendency to falsely recognize novel stimuli that are visually similar to seen stimuli. Visual false memory also occurs when stimuli are meaningful, suggesting that semantic information interferes with the encoding of visual details. However, the neural mechanisms of this semantic interference effect are largely unknown. In the present fMRI study, participants were scanned while encoding visually similar fonts presented with words (word-fonts) or pseudowords (pseudoword-fonts), and later, when recognizing old, new similar (lures), and new dissimilar (novel) fonts displayed in the same meaningless letter string. We performed (1) representational similarity analysis (RSA) at encoding to identify visual, visuosemantic, and semantic representations associated with subsequent visual true and false font recognition, (2) encoding-retrieval similarity (ERS) analysis to assess their reinstatement during retrieval, and (3) mediational analyses to examine hippocampal contributions. The study yielded three main findings. First, visuosemantic representations supported true font recognition when stored in right fusiform gyrus, but false recognition of word-fonts when stored in the left fusiform gyrus. Second, mirroring this pattern, reinstatement in right fusiform gyrus was associated with true font recognition, whereas reinstatement in left fusiform gyrus was linked to false recognition of word-fonts. Finally, posterior hippocampal activation reduced false font memory mainly for pseudoword-associated fonts via decreased reinstatement in perceptual regions, while anterior hippocampal activity increased false memory of word-fonts via enhanced reinstatement in semantic regions. Taken together, these findings reveal how distinct hippocampal-cortical pathways differentially bias memory towards perceptual specificity or semantic generalization.

**Significance Statement:** False memories are often triggered by visual similarity, but this study shows that meaning encoded during learning can distort memory for visual details, even when retrieval cues are meaningless. Participants learned fonts associated with words or pseudowords and judged whether similar lure fonts, shown on a meaningless letter string, were seen before. Although behavioral performance was similar across conditions, brain imaging revealed a key dissociation: the left fusiform gyrus and anterior hippocampus promote semantic generalization that increases false recognition, whereas the right fusiform gyrus and posterior hippocampus support perceptual specificity that protects against it. These findings reveal how distinct hippocampal–cortical pathways differentially bias memory toward truth or illusion.

## Introduction

Despite the remarkable capacity of human visual memory (1, 2), people often falsely recognize new images that are visually similar to previously seen ones. Although this effect can occur for meaningless images (3–6), it is also observed when images are meaningful or are paired with meaningful information (5, 7). This pattern suggests that semantic processing can interfere with the encoding of visual details. Visual false recognition of meaningless stimuli has been associated with reduced activity in posterior brain regions (6), such as early visual cortex (**EVC**) and lingual gyrus (**LG**). However, the neural mechanisms underlying semantic interference with visual detail encoding remain largely unknown. Identifying these mechanisms was the primary goal of the present fMRI study.

Rather than posterior visual areas, such as EVC and LG, the semantic interference on visual detail encoding is likely to involve more anterior visual areas that show greater sensitivity to semantic information, such as the left fusiform gyrus. Evidence from both implicit (8–10) and explicit (11) memory studies indicates that whereas the right fusiform gyrus (**RFG**) is primarily sensitive to visual details, the left fusiform gyrus (**LFG**) is also sensitive to the meaning of information. Thus, while RFG tends to behave like posterior visual areas EVC and LG, LFG behaves more like anterior areas, such as left ventral anterior temporal lobes (**LvATL**) and left ventral prefrontal cortex (**LvPFC**). Thus, for simplicity, we refer to RFG, EVC, LG as (visual) *detail-sensitive regions*, and we refer to LFG, LvATL, and LvPFC as (semantic) *meaning-sensitive regions*.

Visual memory also depends on the engagement of regions critical for memory, such as the hippocampus (12, 13). Unlike the hemispheric asymmetry observed in fusiform cortex, the hippocampus exhibits a posterior-anterior gradient: posterior HC (**pHC**) is more sensitive to visual information, while anterior HC (**aHC**) is more sensitive to semantic information (14, 15). Together, these findings suggest that RFG and pHC primarily support visual true memory, whereas LFG and aHC may contribute to the semantic amplification of visual false memory.

In the current study, participants encoded several sets of similar fonts (e.g. fonts with curly lines, fonts with a horizontal streak; **Fig. 1**), which yielded false recognition of novel but similar fonts (e.g. new font with curly lines, new font with a horizontal streak). Semantic interference was manipulated by displaying fonts using either meaningful words (**Wfonts**) or meaningless pseudowords (**Pfonts**). Although participants were instructed to remember the fonts and ignore the words, the automaticity of reading (16) led us to expect that words meaning would interfere with the encoding of fonts’ visual details. During the recognition test, all fonts were displayed using *the same* meaningless letter string (ENIRSTDAHUL), ensuring that semantic interference occurred only during encoding and not during retrieval.

**Figure 1.**
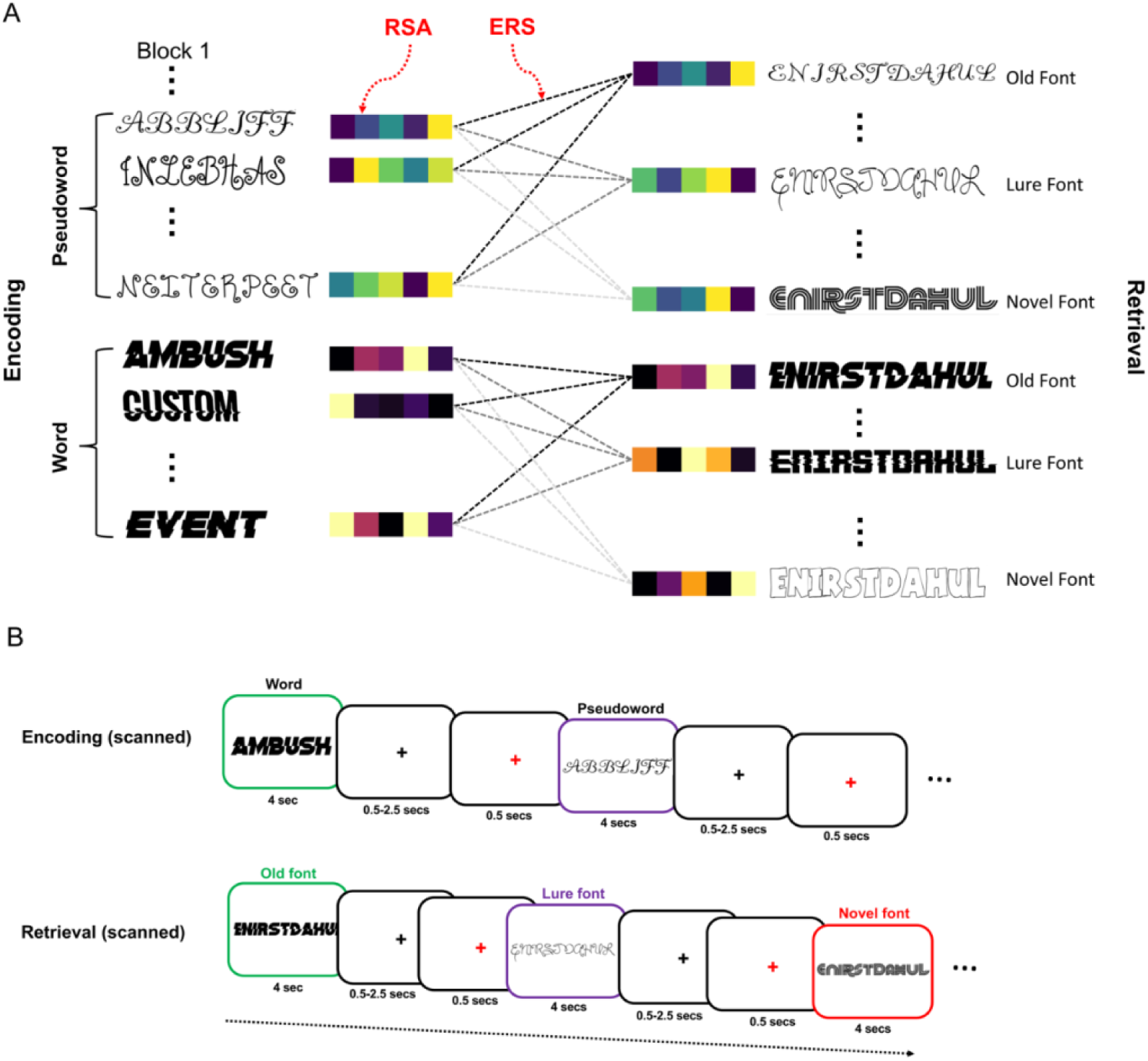
(A) Example of the stimuli. In the example, a set of fonts with curly lines was assigned to pseudowords, and a set of fonts with a horizontal streak was assigned to words. During retrieval, there were for each of these sets, old, lure, and novel fonts. All words in the experiment were presented in German; English translations are shown here for illustrative purposes only (see *Data availability* for the complete stimulus set and norming data). (**B**) **Paradigm**. Separate study and test blocks were used: during encoding blocks, participants rated the pleasantness of each font, and during retrieval blocks they made old/new decisions.

*Representational similarity analyses* (RSA; (17)) were used to measure the strength of visual, visuosemantic, and semantic representations during encoding, which was linked to subsequent true and false recognition of Pfonts and Wfonts. Because memory performance depends not only on the formation of representations during encoding but also on their reinstatement during retrieval, we measured *encoding-retrieval similarity* (13, 18). Finally, we used *mediational analyses* to quantify the effect of HC activation on ERS. Given that effects of encoded representations on subsequent memory depend not only on representation type but also on brain region (19, 20) we expected that both factors would interact with the manipulation of meaning (Wfont vs. Pfont) during encoding.

We tested three predictions. First, for RSA during encoding, we predicted that visual representations in detail-sensitive regions (EVC, LG, and RFG) would boost subsequent true font recognition and attenuate subsequent false font recognition, while semantic representations in meaning-sensitive regions (LFG, LvATL, and LvPFC) would have the opposite effect: reduce subsequent true front recognition and increase subsequent false font recognition. Given their hybrid nature, the effects of visuosemantic representations are more difficult to predict, but it is likely their effects on subsequent memory would depend on the region in which they are encoded. Second, for ERS results, we predicted that greater ERS would be associated with visual true recognition in detail-sensitive regions, reflecting perceptual reinstatement, but with false recognition in meaning-sensitive regions, reflecting semantic reinstatement. Finally, we assumed that pHC activation, which has been related to the encoding of visual features, and aHC activation, which has been linked to the encoding of semantic features, would modulate ERS, impacting memory performance in different ways. Specifically, we predicted that pHC activation would boost ERS in detail-sensitive regions, enhancing true recognition, while aHC activation would boost ERS in meaning-sensitive regions, enhancing false recognition.

## Results

### Memory performance

During encoding, fonts rated as more pleasant had better subsequent visual true recognition, both for Pfonts (b = 0.14, SE = 0.05, z = 2.81, p = 0.005) and Wfonts (b = 0.24, SE = 0.05, z = 4.99, p < 0.001), with no difference between them (b =-0.10, SE = 0.07, z =-1.54, p = 0.123). Encoding reaction times did not differ for Pfonts and Wfonts (b = 0.00, SE = 0.01, t = 0.01, p = 0.991), indicating that differences in fMRI subsequent memory effects between them cannot be attributed to differences in time-on-task.

During retrieval, font recognition was similar for Pfonts and Wfonts (see *SI Appendix*, *Memory performance*, *Fig. S1).* We assessed recognition performance with the Pr index (hits minus false alarms), collapsing across confidence levels due to very few low-confidence responses. Visual true recognition was high for both Pfonts (M = 0.56, SD = 0.12) and Wfonts (M = 0.60, SD = 0.09), with no reliable difference between them (b = 0.03, SE = 0.02, t = 1.76, p = 0.09). Visual false recognition was above chance for both Pfonts (M = 0.34, SD = 0.13) and Wfonts (M = 0.36, SD = 0.11), with no significant difference between them (b = 0.01, SE = 0.02, t = 0.61, p = 0.54).

In sum, there were no significant differences between Pfonts and Wfonts on (1) the effect of pleasantness on subsequent memory, (2) encoding RTs, (3) true recognition, and (4) false recognition. Given the similarity of Pfonts and Wfonts in behavior, fMRI differences between these stimuli cannot be attributed to difficulty differences.

### RSA: Effects of encoded representations on subsequent visual true and false memory

Before testing our prediction regarding the effects of encoded representations on subsequent visual true and false memory, we investigated the distribution of representations across ROIs. The results of these analyses can be found in *SI Appendix*, *RSA analyses independent of subsequent memory* (*Fig. S2* and *Table S1*).

Our first prediction, which was tested with RSA, was that visual representations in detail-sensitive regions (EVC, LG, and RFG) would boost subsequent true font recognition and attenuate subsequent false font recognition, while semantic representations in meaning-sensitive regions (LFG, LvATL, and LvPFC) would have the opposite effect, namely to reduce subsequent true front recognition and increase subsequent false font recognition. Given the hybrid nature of visuosemantic representations we could not make a specific prediction but we expected that their behavior would depend on the region in which they were encoded. The results (purple and teal stars in **Fig. 2**, see also *SI Appendix*, *Table S2*) were largely consistent with our prediction.

**Figure 2.**
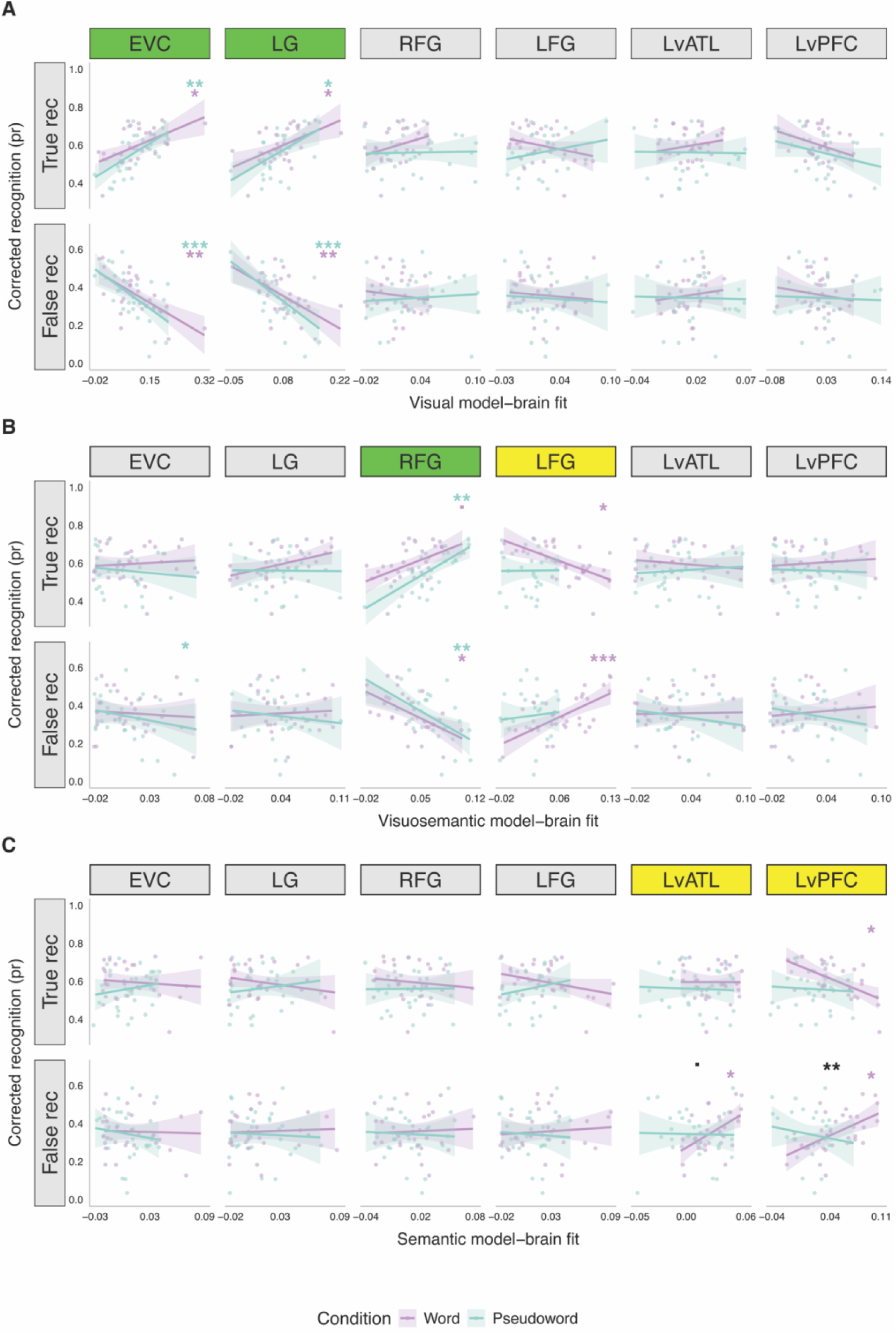
Visual (**A**), visuosemantic (**B**), and (**C**) semantic model–brain fit predict true and false recognition differently for studied fonts. Line plots show corrected hit rate as a function of model-brain similarity, split by encoding condition (Wfonts: purple; Pfonts: teal). Shaded areas indicate SE across subjects. Colored asterisks denote significant linear trends per ROI (FDR-corrected per 6ROIs); black asterisks indicate significant differences in slope between Wfont and Pfont conditions within a ROI; (FDR-corrected for 2 planned contrasts); *p <.05, **p <.01, ***p <.001.

Despite the large number of graphs in **Fig. 2**, there were just two opposing effects (highlighted with green and yellow headings): in detail sensitive regions (EVC, LG, RFG), representational strength predicted *higher* true recognition and *lower* false recognition for both Wfonts and Pfonts, whereas in meaning-sensitive regions (yellow headings: LFG, LvATL, and LvPFC), representational strength was associated with *lower* true memories and *higher* false memories for Wfonts. To investigate these effects in detail, we performed two groups of analyses, one focused on differences between Pfonts and Wfonts, and another focused on differences between true and false recognition.

### Differences between Pfonts and Wfonts

In the general model, the effects of visual or visuosemantic representations on true and false recognition did not differ for Wfonts vs. Pfonts, while the effect of semantic representations in LvPFC on false recognition was significantly greater for Wfonts than Pfonts (black stars in **Fig. 2**, see also *SI Appendix*, *Table S3*). However, an analysis focused on visuosemantic representations in RFG vs. LFG showed that (1) true recognition was greater for Pfonts in RFG > LFG (ß = 2.37, SE = 1.16, z = 2.04, p =.041) and lower for Wfonts in LFG > RFG (ß = - 2.88, SE = 0.83, z = 3.48, p =.001); and that (2) false recognition was lower for Pfonts in RFG > LFG (ß =-3.31, SE = 1.17, z =-2.83, p =.005) and greater for Wfonts in LFG > RFG (ß = 3.68, SE = 0.79, z = 4.66, p <.001). Thus, visuosemantic representations behaved like visual representations when stored in RFG but like semantic representations when stored in LFG.

### Differences between true and false recognition

Finally, we examined planned contrasts between true and false recognition (**Fig.2**, see also *SI Appendix*, *Table S4*). Visual representations in posterior regions (EVC and LG) predicted higher true recognition and lower false recognition for both Wfonts and Pfonts. This effect was significant in EVC for Wfonts (ß = 1.80, SE = 0.45, z = 3.98, p <.001), LG for Wfonts (ß = 2.28, SE = 0.60, z = 3.81, p <.001), EVC for Pfonts (ß = 2.56, SE = 0.49, z = 5.22, p <.001), and LG for Pfonts (ß = 3.30, SE = 0.68, z = 4.84, p <.001). Visuosemantic representations in RFG showed a similar pattern: they predicted higher true recognition and lower false recognition for Wfonts (ß = 3.60, SE = 0.96, z = 3.75, p <.001) and Pfonts (ß = 5.23, SE = 1.06, z = 4.95, p <.001). However, the opposite pattern emerged in LFG, where visuosemantic representations predicted lower true recognition and higher false recognition, selectively for Wfonts (ß =-2.95, SE = 0.63, z = - 4.66, p <.001). Likewise, semantic representations in LvPFC predicted lower true recognition and higher false visual recognition only for Wfonts (ß =-3.19, SE = 0.80, z =-3.97, p <.001).

In sum, mostly consistent with our predictions, RSA yielded three effects. First, visual representations in posterior visual cortex (EVC, LG) were associated with accurate recognition (higher true, lower false) regardless of condition. Second, visuosemantic representations in FG varied with hemisphere and condition: in RFG, they were associated with higher true recognition (only Pfonts) and lower false recognition (both Pfonts and Wfonts), whereas in LFG, they were associated with higher false recognition and lower true recognition (only Wfonts). Finally, semantic representations in anterior brain regions (LvATL, LvPFC) were associated with higher visual false recognition (only Wfonts). In LvPFC, this effect was accompanied by a decrease in true recognition. Thus, cortical representations predict subsequent visual memory depending on (1) their nature (visual representations enhance true memory and semantic, false memory), (2) the location where they are stored (visuosemantic representations promote true memory in RFG but false memory in LFG), and (3) the encoding manipulation (Pfont promote true memory in RFG but false memory in LFG). Possibly due to their hybrid nature, visuosemantic representations in fusiform gyrus were the most sensitive to location and manipulation effects.

#### ERS: Effects of reinstatement on visual true and false memory

The effects of representations on memory depend not only on the storage of visual, visuosemantic, and semantic representations during encoding, which we measured with RSA, but also on the reactivation of these representations during retrieval, which we measured with ERS. We linked ERS to true recognition by comparing Hit vs. Miss, and to false recognition by comparing FA vs. CR. We then determined whether these effects differed by Stimulus Type (Wfonts vs. Pfonts). Results are shown below in **Fig. 3** and reported fully in *SI Appendix*, *Table S5* (both within-and across condition ERS values).

**Figure 3.**
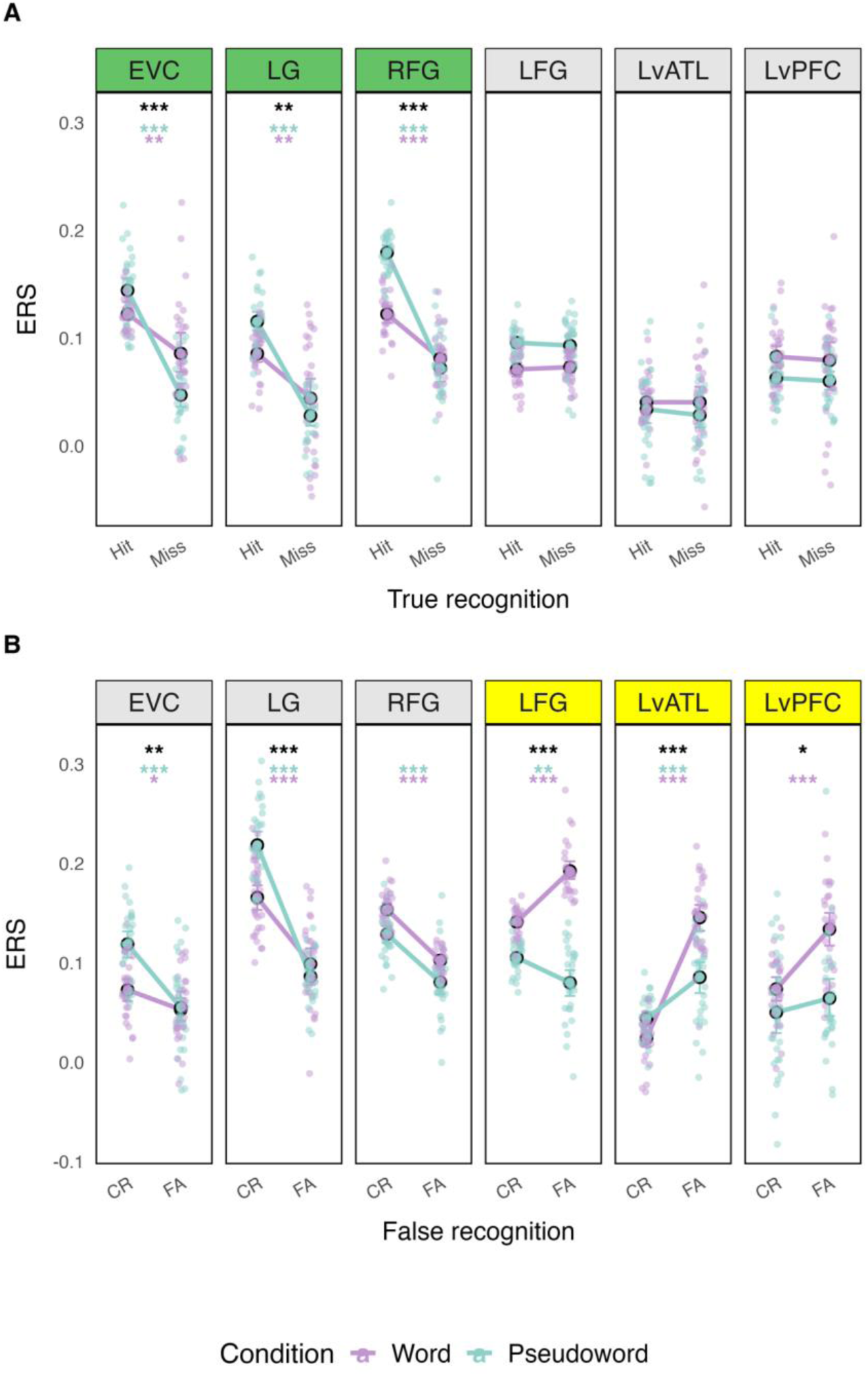
Brain regions showing significant associations between ERS and memory. Line plots display ERS values for (**A**) true recognition condition (Hit vs. Miss) and (**B**) false recognition condition (FA vs. CR). Error bars represent the standard error of the mean across participants. Colored asterisks above the lines indicate p-values for paired t-tests comparing memory outcomes within each condition (Hit ≠ Miss, FA ≠ CR). Black asterisks indicate p-values for paired t-tests comparing stimulus type (Word vs. Pseudoword) within each memory contrast (Hit ≠ Misses, FA ≠ CR). All p-values are FDR-corrected across the six ROIs. Significance thresholds: *p < 0.05, **p < 0.01, ***p < 0.001.

For ERS results, we predicted that greater ERS would be associated with visual true recognition in detail-sensitive regions (EVC, LG, and RFG), reflecting perceptual reinstatement, but with visual false recognition in meaning-sensitive regions (LFG, LvATL, and LvPFC), reflecting semantic reinstatement. ERS results were generally consistent with these predictions. As shown in **Fig. 3**, in detail-sensitive regions, ERS was associated with higher true recognition (Hit > Miss) for both Wfonts and Pfonts (green headings), while in meaning-sensitive regions, ERS was linked to higher false recognition (FA > CR) for Wfonts (yellow headings). Turning to direct Pfont vs. Wfont contrasts (black asterisks), true recognition was significantly greater for Pfonts than Wfonts in all detail-sensitive regions, and correct rejection was greater for Pfonts than Wfonts in EVC and LG. In meaning-sensitive regions, false recognition was reliably greater for Wfonts than Pfonts.

To investigate RFG-LFG dissociation, we conducted within-subject paired t-tests comparing memory-related contrasts between the two regions. These analyses showed that the Hit > Miss difference was larger in RFG than LFG for both Wfonts (t(29) = 4.59, p <.001, d = 1.37) and Pfonts (t(29) = 11.73, p <.001, d = 3.09), suggesting that RFG differently supports true recognition independently of stimulus type. Conversely, the FA > CR difference was larger in LFG than RFG for Wfonts (t(29) = 14.82, p <.001, d = 3.88) but not for Pfonts (t(29) = 1.95, p <.061, d = 3.09). Thus, the role of LFG in false recognition depended on the presence of semantic information during encoding.

#### Mediation analysis: Hippocampal contributions to memory via pattern similarity

Finally, our third prediction was that the hippocampus would modulate the reinstatement of visual and semantic features. Specifically, we predicted that greater pHC activation would enhance ERS associated with true recognition in detail-sensitive regions (EVC, LG, and RFG), while aHC activation would increase ERS linked to false recognition in meaning-sensitive regions (LFG, LvATL, and LvPFC). The results were largely consistent with this prediction in all ROIs (see *SI Appendix*, *Tables S6* and *S7*). **Fig. 4** focuses on RFG and LFG, as representative ROIs where perceptual and semantic influences diverged. ERS in RFG supported true recognition of Pfonts, as well as mnemonic discrimination of similar lures, while ERS in LFG contributed to false recognition of Wfonts.

**Figure 4.**
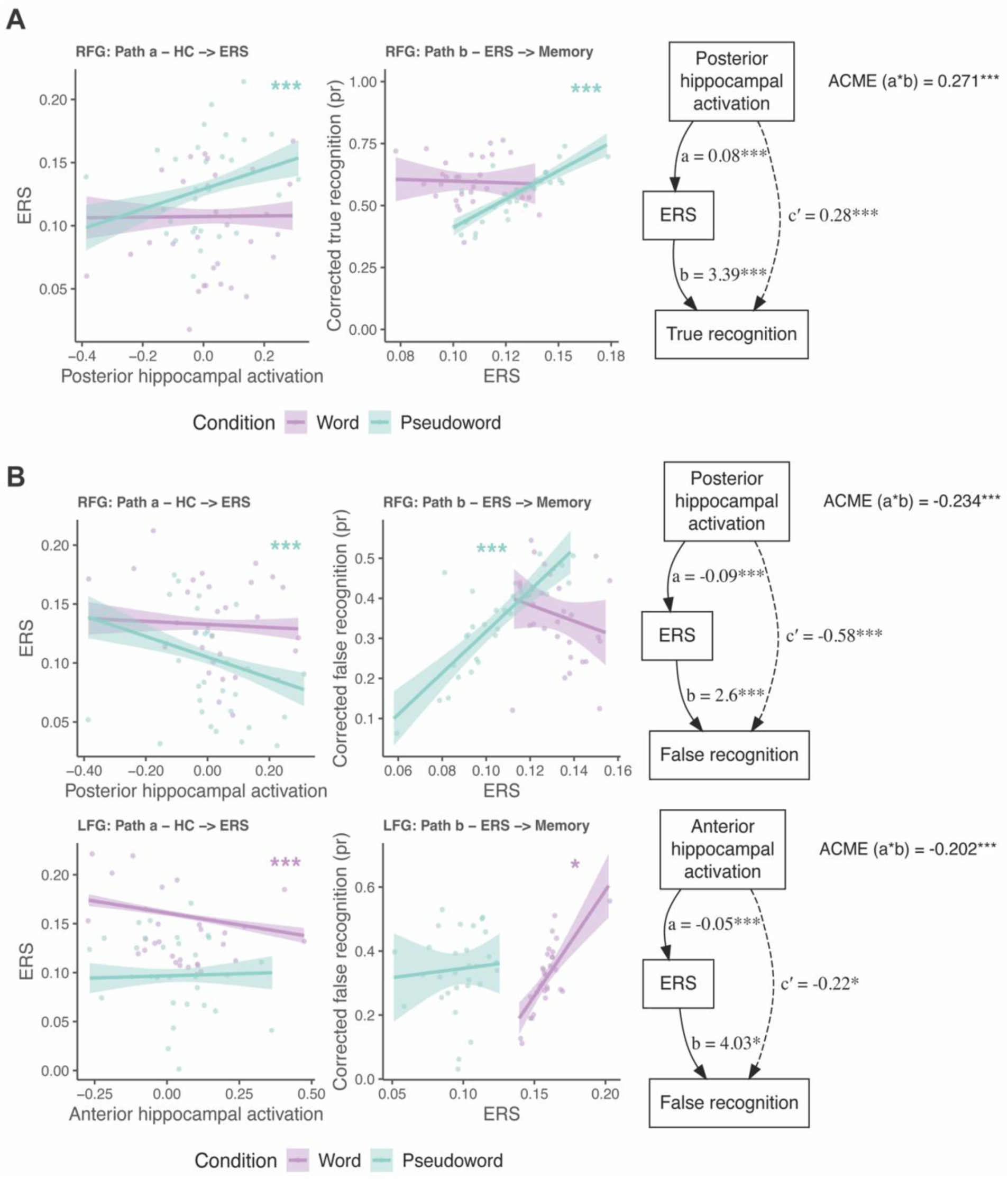
ERS in RFG and LFG mediates the link between hippocampal activation and false recognition. (**A**) Posterior hippocampus: the positive association with true recognition of fonts was partially mediated by RFG ERS. (**B**) Posterior and anterior hippocampus: the negative association with false recognition of similar lure fonts was also partially mediated by FG ERS. *P*-values reflect FDR-corrected mediation significance across ROIs. *p* <.05, p <.01, *p* <.001.

Consistent with expectations, pHC activation was associated with higher true recognition via increased ERS in EVC (ACME = 0.21, 95% CI [0.07, 0.35], p =.006) and RFG (see **Fig. 4A**; ACME = 0.27, 95% CI [0.12, 0.43], p <.001) only for Pfonts. Also, pHC activation reduced false recognition via greater ERS in EVC (ACME =-0.16, 95% CI [-0.28,-0.03], p =.012) and RFG (see **Fig. 4B**, upper panel; ACME =-0.23, 95% CI [-0.35,-0.11], p <.001), again only for Pfonts. Finally, aHC activation boosted false recognition via ERS in LFG (see **Fig. 4B**, lower panel; ACME =-0.20, 95% CI [-0.35, - 0.04], p =.030) and LvATL (ACME =-0.18, 95% CI [-0.34,-0.02], p =.030) but only for words.

Together, these findings indicate that HC subregions differentially modulate cortical reinstatement based on stimulus type and memory outcome. pHC enhanced perceptual reinstatement during true recognition of Pfonts but suppressed reinstatement during lures, supporting discrimination. In contrast, aHC increased reinstatement in anterior regions during false recognition of Wfonts, suggesting inappropriate semantic reinstatement. This double dissociation suggests that semantic context can disrupt perceptual memory by triggering non-diagnostic reinstatement, whereas its absence supports more selective retrieval.

## Discussion

This study reveals how semantic information, present only during encoding, can reshape perceptual memory for fonts and lead to recognition errors. Using a multivariate approach that combines RSA, ERS, and mediation by HC activation, we demonstrate that semantic information is not only encoded alongside perceptual details, but it can be reactivated in ways that distort memory. This semantic reinstatement occurred in a task with no meaningful cues at test, underscoring the automaticity and potency of semantic interference during encoding.

Our findings support a constructive view of memory in which true and false recognition arise from different balances of perceptual specificity and semantic generalization. The presence of a word at encoding, despite the instruction to disregard it and being irrelevant at retrieval, was sufficient to trigger semantic reinstatement, biasing recognition in mid (LFG) and anterior (LvATL) ventral pathway regions. In other words, semantic context encoded early can infiltrate and distort later mnemonic decisions—even when those decisions should, in theory, rely solely on visual cues.

The results align with theoretical accounts such as fuzzy-trace theory (21, 22), which posits that memory stores both perceptual (verbatim) and semantic (gist) traces, and that gist-based retrieval during lures yields false recognition. Our data suggest that the gist trace, here, the semantic meaning of a word, was automatically activated at test and overrode the perceptual trace in specific cortical regions. Dual-code theory (23) similarly predicts such interference when verbal and visual codes are processed in parallel. Our findings give neurobiological grounding to these cognitive theories by identifying the circuits through which perceptual and semantic information compete or converge.

RSA at encoding showed that semantic information, especially that captured by SGPT (a language model based on distributional semantics), was more strongly represented for fonts associated with words (Wfonts) than for fonts associated with pseudowords (Pfonts). In contrast, visual representations were localized to posterior regions, such as early visual cortex (EVC) and lingual gyrus (LG), while visuosemantic representations emerged along the ventral stream, peaking in FG. This posterior-to-anterior representational gradient mirrors established models of visual and semantic integration (24, 25). Notably the LvATL, a putative transmodal semantic hub, showed heightened representation of semantic features for Wfonts, despite the task’s purely visual demands. Critically, these semantic traces were not just encoded, but they were recapitulated during retrieval. ERS analyses showed that anterior regions reinstated encoding patterns more strongly for Wfonts than Pfonts, particularly for false alarms. This suggests that semantic context modulated both encoding and retrieval dynamics.

The fusiform gyrus (FG) stood out as a functional intersection between perception and meaning. It showed strong RSA effects for both visual and visuosemantic models, especially CLIP, which fuses image and text data. In FG, CLIP-based model-brain fit predicted increased false recognition for words, but decreased false alarms for pseudowords. A similar pattern emerged for AlexNet’s layer 2, a purely visual model, where greater representational specificity in FG and EVC was associated with reduced memory errors. These double dissociations suggest that FG does not merely reflect visual similarity but flexibly integrates perceptual and semantic input depending on context. When fonts are encoded in the context of words (Wfonts), semantic generalization may dilute perceptual precision. In contrast, when words are encoded in pseudowords (Pfonts), reliance on visual features may boost mnemonic accuracy. FG biases recognition toward or away from false recognition depending on the encoding context.

LvATL also emerged as a driver of semantic reinstatement during false memory. It exhibited increased ERS for false alarms to word-associated fonts and was specifically modulated by anterior hippocampal activation. These findings echo prior work identifying vATL as a transmodal semantic region involved in conceptual abstraction (25, 26). In our case, it appears to have generalized from previously encoded semantic input and misapplied that information at retrieval—suggesting that false memories can arise not from confusion between similar perceptual inputs, but from semantically driven misattribution.

The hippocampus played a central, but bifurcated, role. Posterior hippocampal (pHC) activation reduced false memory via weaker ERS in early visual and right fusiform regions, consistent with its role in perceptual precision and pattern separation (15, 27). In contrast, anterior hippocampal (aHC) activation increased false recognition for Wfonts via stronger ERS in LvATL and LFG. This anterior-posterior dissociation supports theories of hippocampal functional specialization, where the posterior segment supports detailed encoding and the anterior segment facilitates integration and semantic generalization (14, 28). Importantly, mediation analyses confirmed this directional pattern: pHC suppressed false memory through reduced ERS in perceptual regions, while aHC amplified it through enhanced ERS in semantic hubs, but only for stimuli with encoded meaning (Wfonts). These findings point to a dual mechanism through which the hippocampus regulates the content and outcome of memory reinstatement—not simply whether a trace is reactivated, but what kind of trace is reinstated and to what effect.

Taken together, these neural dynamics underscore the constructive nature of episodic memory (29). The semantic influences we observed were not triggered by external cues but emerged internally from prior reading. The fact that meaningless test strings elicited differential reinstatement patterns based on prior semantic context challenges the assumption that perceptual recognition can be cleanly isolated from meaning. In real-world settings, where perception and meaning are inextricably linked, such interactions likely occur even more frequently and may underlie both adaptive generalization and harmful memory distortions.

Direct comparisons of true versus false memory also support a representational dissociation. While both forms of recognition involved reinstatement, the nature of the reactivated trace differed: perceptual reinstatement supported true memory, particularly for pseudowords; semantic reinstatement increased false recognition, particularly for words. FG again showed a reversal: visuosemantic RSA in FG enhanced true recognition for pseudowords but impaired recognition for words. These findings argue against a unidimensional memory strength model and instead support a multidimensional view where memory accuracy reflects the balance between perceptual fidelity and semantic intrusion (30, 31).

In sum, our findings reveal a cortical–hippocampal network that transforms visual input into increasingly abstract representations, which are later reactivated to support, or distort, memory. Early visual areas encode precise perceptual details. FG integrates these with semantic information. LvATL abstracts meaning across episodes. The hippocampus modulates which trace is prioritized at retrieval: perceptual precision or semantic generalization. False memories, therefore, are not merely accidents of confusion but predictable outcomes of how distributed systems interact across representational hierarchies. Future work should examine how attentional states or task goals at encoding modulate these dynamics, and how this system may become unbalanced in aging or clinical populations. The spontaneous reinstatement of semantic content, even in perceptual tasks, may be a key contributor to real-world memory errors, and a target for intervention.

## Materials and Methods

### Participants

Sample size (N = 30) was determined based on effect sizes reported in previous studies of RSA, ERS, and subsequent memory (20, 32, 33). The participants were 33 right-handed, young native German speakers, with normal or corrected vision, and no psychiatric, neurological, or major medical history. Three were excluded due to poor memory performance, resulting in a final sample of 30 participants (M = 23.97 years, SD = 4.32; 23 females). Older participants were also tested but their results will be reported elsewhere. All gave written informed consent to a protocol approved by Humboldt University Ethics Committee (Ref. 2022-05), and received compensation or course credit.

### Stimuli

There were three types of stimuli: (1) 84 abstract German words (34), with a length of 5–11 letters, a familiarity > 2.19, and abstractness > 2.19; (2) 84 pseudowords generated with *Wuggy* (35) and matched to words on orthographic and phonological properties; and (3) 441 fonts belonging to 21 distinct sets, such as fonts with curly lines (21 fonts each). The experiment consisted of 7 study–test blocks, and each block included two 21-font sets, one displayed as words, and one as pseudowords (**Fig. 1A**). For each of these 21-font sets (e.g., curly fonts), 12 fonts were presented during encoding and represented in the retrieval list (*old fonts*). For each set, the retrieval list included the remaining 9 fonts of the 21-font set (*lure fonts*) and 9 fonts from a separate dissimilar font set (*novel fonts*). Stimulus–condition assignments and trial order were randomized across participants. Eighteen filler trials preceded the first block.

### Experimental Design

The fMRI session consisted of 14 runs, comprising 7 study–test blocks. During encoding, participants viewed each font, and rated how much they liked the font (not the word) from 1 = “not at all” to 4 = “very much.” Each font appeared for 4 s, and was followed by a jittered fixation (mean SOA = 6 s). Participants were informed that their memory for the fonts (not the words) would be tested. During retrieval, old, lure, or novel fonts were displayed with the same string of 11 most frequent German letters (“ENIRSTDAHUL”), and participants made an old/new decision with confidence judgments. Confidence responses were collapsed and are not reported or analyzed here. Response-hand mappings were counterbalanced across participants.

### fMRI Acquisition and Preprocessing

Imaging data were acquired with a Siemens 3T Prisma scanner with a 64-channel head coil. Functional images were collected using a multiband T2*-weighted EPI sequence (TR = 800 ms, TE = 37 ms, flip angle = 52°, 2 mm³ voxels, 72 slices, FoV = 208 mm). A high-resolution T1-weighted structural scan (MPRAGE; 0.8 mm³) and spin echo field maps were also acquired. Data preprocessing was performed using *fMRIPrep* version 23.0.1 (36), which is based on *Nipype* 1.8.5 (37). Unless otherwise stated, all parameters followed default settings. For full preprocessing details, see *SI Appendix, Image preprocessing*.

### Regions of Interest (ROIs)

Based on prior work on memory and visual or semantic processing (19, 20, 38), 8 ROIs were defined using three atlases. EVC was defined using probabilistic cytoarchitectonic maps from the Jülich atlas (39). LG, RFG, LFG, LvATL, and LvPFC [BA44/45] were from the Harvard–Oxford structural atlas (40). Finally, pHC and aHC were from the Brainnetome atlas (Fan et al., 2016). ROIs were defined in MNI space and transformed into each subject’s native space using inverse deformation fields from *fMRIPrep* (see *SI Appendix*, *ROI*, *Fig. S3*).

### Behavioral analyses

We used a generalized linear mixed-effects model (GLMM, R Core Team, 2023) to test whether pleasantness ratings (from 1 to 4) predicted subsequent memory (hit vs. miss). Encoding response times (RTs) were analyzed with a linear mixed-effects model (LMM) with Stimulus Type (word vs. pseudoword) as predictor. Recognition accuracy (Pr = hit − false alarm rate) was compared across conditions using linear mixed-effect models.

### RSA analyses independent of subsequent memory

Before RSA, we estimated one beta image per encoding trial using a least-squares-all (LSA) approach implemented in SPM12 (41). Each participant’s design matrix included a separate regressor per trial, convolved with a canonical hemodynamic response function. Each run also included six motion regressors and a session constant. Models were fit to preprocessed native-space images using an AR(1) autocorrelation model with a 128-second high-pass filter. Data were normalized to a grand mean of 100. A brain mask included voxels with at least 10% probability of gray or white matter based on T1 segmentation.

RSA was performed using standard procedures (17). Model representational dissimilarity matrices (RDMs) were computed separately for word and pseudoword conditions based on 84 font stimuli. The visual RDM captured low-level image features using layer 2 of AlexNet (42) via the `ThingsVision` toolbox (43). Visuosemantic RDMs were based on CLIP embeddings (44), and semantic RDMs used SGPT sentence embeddings (45). For visual and visuosemantic models, dissimilarity was defined as 1 minus the Pearson correlation between feature vectors; for semantic models, as 1 minus the cosine similarity between text-based embeddings. Each RDM was a symmetric 84×84 matrix, with higher values indicating greater predicted dissimilarity.

Brain RDMs were computed for each region of interest (ROI) and participant by calculating 1 minus the Pearson correlation between multivoxel activity patterns for all pairs of encoding trials. For each model, the second-order model–brain fit was quantified by correlating (Spearman correlation) the lower triangle of the model RDM with the corresponding brain RDM, separately for each ROI, subject, and stimulus type. This yielded one similarity score per Representation Type × ROI × Stimulus Type × Subject.

Fisher-transformed model–brain similarity scores were analyzed using one-sample and paired-samples t-tests, as well as repeated-measures ANOVA, depending on the number of within-subject factors. One-sample and paired-samples t-tests (Wfonts vs. Pfonts) were performed within each model and ROI to test for (1) above-chance similarity and (2) stimulus-type differences (see *SI Appendix*, *RSA analyses independent of subsequent memory*). Reported p-values were corrected using the false discovery rate (FDR) method across ROIs and Stimulus Type. In the main analysis, we tested group-level differences across ROI categories using paired t-tests (e.g., early vs. late visual regions for AlexNet, semantic vs. nonsemantic regions for SGPT). To examine lateralization and content sensitivity, repeated-measures ANOVAs assessed Representation Type × ROI × Stimulus Type effects in FG and in semantic ROIs. All tests were conducted in R using the `afex`, `effsize`, and `BayesFactor` packages.

### RSA analyses predicting subsequent memory

To examine how RSA at encoding relates to memory performance, we fit a beta regression model predicting corrected recognition (pr) as a function of ROI, Stimulus Type (Wfonts vs. Pfonts), Memory Condition (true vs. false recognition), Representation Type (visual, visuosemantic, semantic), and model–brain fit, including all interactions. Model fit was computed at encoding using representational similarity analysis (RSA) between brain RDMs and model RDMs (see RSA analyses independent of subsequent memory). We estimated the effect of model–brain fit (slope) for each ROI × Memory Condition × Stimulus Type × Representation Type combination. False discovery rate (FDR) correction was applied across 6 ROIs. We also tested 4 planned contrasts across Stimulus Type and Memory Condition using estimated marginal trends. To assess lateralization, we refit the model using only the visuosemantic representation and compared the slope of model–brain fit between LFG and RFG separately for each combination of Stimulus Type (Wfonts, Pfonts) and Memory Condition (true, false recognition). Reported p-values were corrected using the false discovery rate (FDR) method across ROIs and Stimulus Type.

### Encoding-retrieval neural pattern similarity analysis (ERS)

For each ROI, we computed ERS by correlating voxelwise activity patterns from encoding and retrieval trials (Hit, Miss, FA, and CR) using single-trial beta images. ERS was defined as the mean Fisher z-transformed Pearson correlation between each retrieval item and its corresponding encoding item within the same block and Stimulus Type (i.e., Wfonts, Pfonts; see **Fig. 1A**). To isolate item-specific similarity related to font identity, we subtracted a baseline: the average similarity between each retrieval item and encoding items from different font families. Resulting baseline-corrected ERS values were averaged across blocks. Paired t-tests assessed memory effects and word/pseudoword differences. All p-values are FDR-corrected across the six ROIs (46).

### Modulatory effect of hippocampal activity on ERS

To assess how univariate hippocampal activation during encoding modulates ERS-measured reinstatement (13), we conducted mediation analyses. For each subject, we computed the average encoding-phase activation within posterior and anterior HC, separately for Wfonts and Pfonts. We then tested whether hippocampal activation predicted both ERS and subsequent memory outcomes (true and false recognition), and whether ERS mediated these effects. In the mediation model, X was hippocampal activation, M was ERS, and Y was memory performance. Indirect effects were estimated using a quasi-Bayesian Monte Carlo approximation as implemented in the `mediation` package (47).

### Data availability

All code, stimuli (words, pseudowords, fonts), norming variables, and analysis materials will be made available in an open repository upon publication.

## Acknowledgments and funding sources

The study was supported by the Einstein Foundation Berlin (EPP-2017-423) and National Institute of Health (NIA 1RF1AG066901). We thank Paola Gega, Mindia Winkert, Yasamin Azizi, Teuta Dzaferi, and Diana Paola Americano Guerrero for their substantial assistance with data collection.

## Notes

### Competing Interest Statement

The authors have declared no competing interest.

