## Supporting Information for "When knowledge interferes with perception: Neural mechanisms of the semantic amplification of visual false memory"

### Results

#### *Memory performance*

See behavioral results below in Fig. S1.

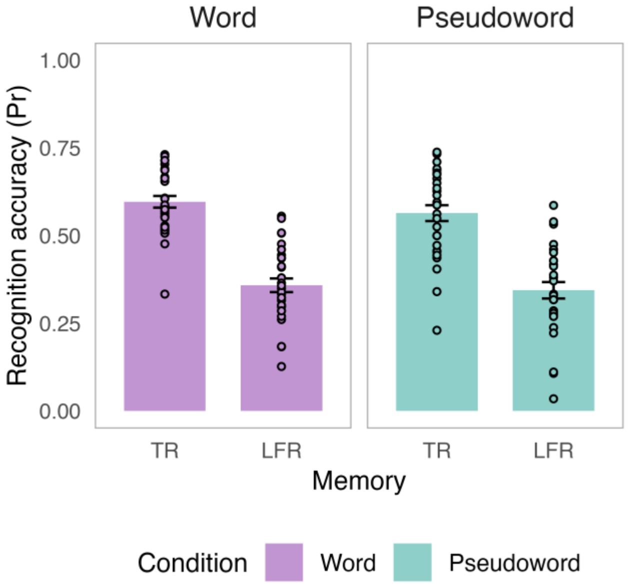


**Figure S1**. Corrected true and false recognition of Wfonts and Pfonts. Bar plot of memory performance, as indexed by pr, are shown for both Wfonts (purple) and Pfonts (teal) condition. Error bars indicate across-participant SE. Black asterisks above the bars indicate p values for tests of whether each individual paired t-test for differences between condition (Wfonts, Pfonts) is > 0. TR = True recognition; LFR = Lure false recognition. $*p < 0.05$, $**p < 0.01$, $***p < 0.001$.

#### *RSA analyses independent of subsequent memory*

The aim of the RSA analyses was to investigate the effects of cortical representations on subsequent memory. However, before turning to this goal, we investigated the distribution of visual, visuosemantic, and semantic representations along the ventral pathway and the hemispheric asymmetry of the fusiform gyrus by examining RSA results independent of memory.

Results from one-sample and paired t-tests for each Representation Type and ROI (i.e., above-chance similarity, Wfonts vs. Pfonts) are displayed below in Figure S2 and reported in Table S1.

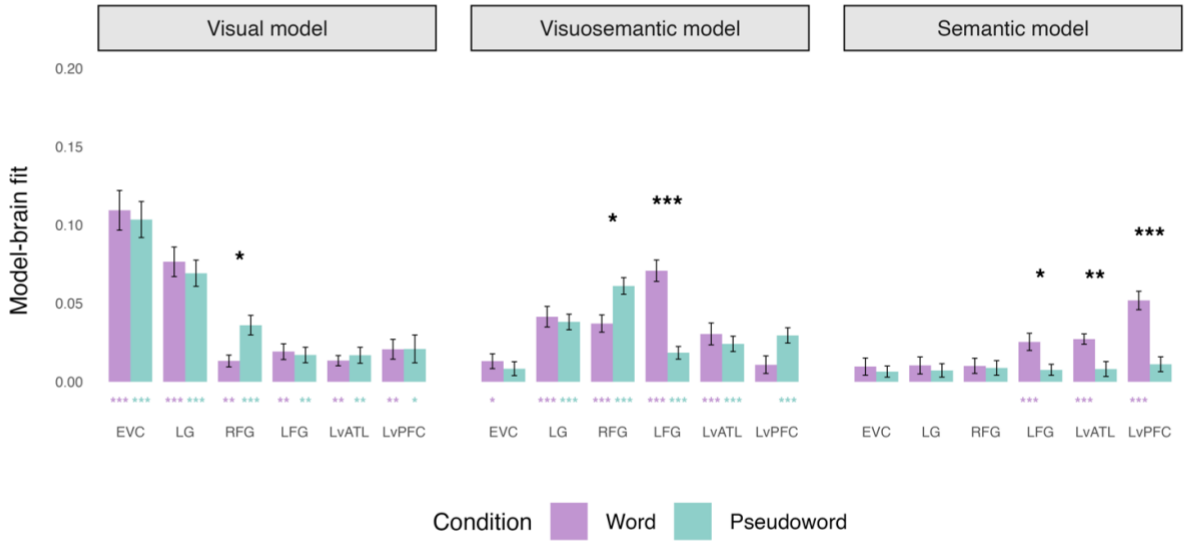


**Figure S2.** RSA results independent of memory showed that visual, visuosemantic, and semantic representations were organized along a posterior-anterior ventral pathway gradient. Bars show mean Spearman correlation between model-based similarity and neural patterns across ROIs. Colored asterisks indicate model–brain fit > 0 (one-sample t-tests); black asterisks mark Wfonts vs. Pfonts differences (paired t-tests). Error bars reflect SEM across subjects. P-values were FDR-corrected across six ROIs: *p < .05, **p < .01, ***p < .001.

Consistent with prior research (1–4), visual representations were significantly stronger in detail-oriented regions (EVC, LG, RFG) than in anterior regions (LvATL, LvPFC, and LFG ($t(29) = 10.43$, $p < .001$, $d = 1.66$)). Interestingly, RFG showed greater visuosemantic coding for Pfonts, and LFG for Wfonts (Representation Type x ROI x Stimulus Type interaction: ($F(1.80, 52.12) = 41.33$, $p < .001$). Finally, semantic representations in LFG, LvATL, and LvPFC were stronger for Wfonts than Pfonts (ROI x Stimulus Type interaction: ($F(1.71, 49.59) = 6.88$, $p = .004$), and stronger than semantic representations in EVC, LG, and RFG ($t(29) = 5.63$, $p < .001$, $d = 0.84$).

**Table S1.** Summary of statistical results for visual, visuosemantic, and semantic representations across six ROIs. For each condition (Wfonts, Pfonts), the table shows one-sample t-tests against zero. Paired t-tests compare representation strength between Wfonts and Pfonts conditions. All tests were performed across participants (n = 30).

| **Visual model** | | | | | | | |
|---|---|---|---|---|---|---|---|
| ROI | Condition | t(29) | p | d | Contrast | p | d |
| EVC | Word | 8.63 | < .001 | 1.58 | 0.44 | .917 | 0.09 |
| | Pseudoword | 8.98 | < .001 | 1.64 | | | |
| LG | Word | 8.13 | < .001 | 1.48 | 0.82 | .917 | 0.15 |
| | Pseudoword | 8.28 | < .001 | 1.51 | | | |
| RFG | Word | 3.52 | .002 | 0.64 | -2.95 | .037 | -0.82 |
| | Pseudoword | 5.77 | < .001 | 1.05 | | | |
| LFG | Word | 3.82 | .001 | 0.70 | 0.30 | .917 | 0.08 |
| | Pseudoword | 3.46 | .003 | 0.63 | | | |
| LvATL | Word | 4.09 | .001 | 0.75 | -0.54 | .917 | -0.15 |
| | Pseudoword | 3.32 | .003 | 0.61 | | | |
| LvPFC | Word | 3.27 | .003 | 0.60 | -0.02 | .984 | -0.01 |
| | Pseudoword | 2.36 | .025 | 0.43 | | | |
| **Visuosemantic model** | | | | | | | |
| ROI | Condition | t(29) | p | d | Contrast | p | d |
| EVC | Word | 2.76 | .012 | 0.50 | 0.70 | .590 | 0.19 |
| | Pseudoword | 1.87 | .072 | 0.34 | | | |
| LG | Word | 6.32 | < .001 | 1.15 | 0.38 | .704 | 0.11 |
| | Pseudoword | 7.66 | < .001 | 1.40 | | | |
| RFG | Word | 6.75 | < .001 | 1.23 | -3.15 | .011 | -0.81 |
| | Pseudoword | 11.51 | < .001 | 2.10 | | | |
| LFG | Word | 10.38 | < .001 | 1.90 | 6.39 | < .001 | 1.72 |
| | Pseudoword | 4.57 | < .001 | 0.84 | | | |
| LvATL | Word | 4.36 | < .001 | 0.80 | 0.71 | .590 | 0.19 |
| | Pseudoword | 4.92 | < .001 | 0.90 | | | |
| LvPFC | Word | 1.94 | .063 | 0.35 | -2.24 | .066 | -0.65 |
| | Pseudoword | 6.03 | < .001 | 1.10 | | | |
| **Semantic model** | | | | | | | |
| ROI | Condition | t(29) | p | d | Contrast | p | d |
| EVC | Word | 1.74 | .092 | 0.32 | 0.53 | .722 | 0.12 |
| | Pseudoword | 1.82 | .104 | 0.33 | | | |
| LG | Word | 1.90 | .081 | 0.35 | 0.64 | .722 | 0.12 |
| | Pseudoword | 1.68 | .104 | 0.31 | | | |
| RFG | Word | 2.08 | .070 | 0.38 | 0.19 | .847 | 0.05 |
| | Pseudoword | 1.88 | .104 | 0.34 | | | |
| LFG | Word | 4.63 | < .001 | 0.85 | 2.44 | .042 | 0.71 |
| | Pseudoword | 2.15 | .104 | 0.39 | | | |
| LvATL | Word | 8.30 | < .001 | 1.51 | 3.40 | .006 | 0.85 |

| | | | | | | | |
|---|---|---|---|---|---|---|---|
| | Pseudoword | 1.70 | .104 | 0.31 | | | |
| LvPFC | Word | 8.78 | < .001 | 1.60 | 6.35 | <.001 | 1.38 |
| | Pseudoword | 2.36 | .104 | 0.43 | | | |

*Effects of encoding RSA on subsequent true and false visual memory*

**Table S2**. Estimated marginal means for the visual, visuosemantic, and semantic models predicting subsequent true and false visual recognition

| Memory | ROI | ß | SE | z-value | p |
|---|---|---|---|---|---|
| | **AlexNet - Visual model** | | | | |
| | EVC | | | | |
| True rec | Word | 0.73 | 0.31 | 2.34 | .033 |
| | Pseudoword | 1.03 | 0.33 | 3.16 | .005 |
| False rec | Word | -1.08 | 0.33 | -3.26 | .004 |
| | Pseudoword | -1.52 | 0.37 | -4.18 | <.001 |
| | LG | | | | |
| True rec | Word | 0.94 | 0.41 | 2.27 | .035 |
| | Pseudoword | 1.23 | 0.46 | 2.68 | .015 |
| False rec | Word | -1.34 | 0.43 | -3.10 | .005 |
| | Pseudoword | -2.07 | 0.50 | -4.11 | <.001 |
| | RFG | | | | |
| True rec | Word | 1.34 | 0.94 | 1.44 | .202 |
| | Pseudoword | 0.06 | 0.59 | 0.11 | .995 |
| False rec | Word | -0.62 | 0.92 | -0.67 | .601 |
| | Pseudoword | 0.00 | 0.56 | 0.01 | .995 |
| | LFG | | | | |
| True rec | Word | -0.89 | 0.74 | -1.20 | .730 |
| | Pseudoword | 0.85 | 0.75 | 1.14 | .730 |
| False rec | Word | -0.42 | 0.70 | -0.60 | .730 |
| | Pseudoword | -0.33 | 0.70 | -0.47 | .770 |
| | LvATL | | | | |
| True rec | Word | 0.79 | 1.09 | 0.72 | .730 |
| | Pseudoword | -0.09 | 0.73 | -0.13 | .899 |
| False rec | Word | 0.74 | 1.12 | 0.66 | .730 |
| | Pseudoword | -0.20 | 0.68 | -0.29 | .844 |
| | LvPFC | | | | |
| True rec | Word | -0.84 | 0.59 | -1.43 | .730 |
| | Pseudoword | -0.63 | 0.42 | -1.49 | .730 |
| False rec | Word | -0.43 | 0.56 | -0.78 | .730 |
| | Pseudoword | -0.27 | 0.39 | -0.69 | .730 |
| | **CLIP - Visuosemantic model** | | | | |
| | EVC | | | | |
| True rec | Word | 0.30 | 0.77 | 0.40 | .754 |
| | Pseudoword | -0.56 | 0.84 | -0.67 | .708 |

| | | | | | |
|---|---|---|---|---|---|
| False rec | Word | -0.44 | 0.74 | -0.59 | .708 |
| | Pseudoword | -1.88 | 0.69 | -2.73 | .019 |
| | LG | | | | |
| True rec | Word | 0.99 | 0.57 | 1.73 | .166 |
| | Pseudoword | -0.05 | 0.74 | -0.07 | .945 |
| False rec | Word | 0.29 | 0.54 | 0.54 | .708 |
| | Pseudoword | -0.48 | 0.71 | -0.67 | .708 |
| | RFG | | | | |
| True rec | Word | 1.57 | 0.68 | 2.31 | .051 |
| | Pseudoword | 2.38 | 0.72 | 3.31 | .006 |
| False rec | Word | -2.03 | 0.68 | -3.00 | .011 |
| | Pseudoword | -2.86 | 0.78 | -3.68 | .003 |
| | LFG | | | | |
| True rec | Word | -1.31 | 0.48 | -2.73 | .038 |
| | Pseudoword | 0.01 | 0.92 | 0.01 | .995 |
| False rec | Word | 1.64 | 0.41 | 3.96 | <.001 |
| | Pseudoword | 0.46 | 0.88 | 0.52 | .926 |
| | LvATL | | | | |
| True rec | Word | -0.38 | 0.52 | -0.73 | .926 |
| | Pseudoword | 0.30 | 0.75 | 0.39 | .926 |
| False rec | Word | 0.09 | 0.51 | 0.17 | .943 |
| | Pseudoword | -1.21 | 0.71 | -1.71 | .264 |
| | LvPFC | | | | |
| True rec | Word | 0.29 | 0.65 | 0.44 | .926 |
| | Pseudoword | -0.16 | 0.76 | -0.21 | .943 |
| False rec | Word | 0.47 | 0.64 | 0.74 | .926 |
| | Pseudoword | -1.25 | 0.71 | -1.75 | .264 |
| | **SGPT - Semantic model** | | | | |
| | EVC | | | | |
| True rec | Word | -0.29 | 0.67 | -0.44 | .937 |
| | Pseudoword | 0.77 | 1.02 | 0.75 | .937 |
| False rec | Word | -0.24 | 0.64 | -0.37 | .937 |
| | Pseudoword | -0.76 | 0.93 | -0.82 | .937 |
| | LG | | | | |
| True rec | Word | -0.70 | 0.68 | -1.03 | .937 |
| | Pseudoword | 0.64 | 0.85 | 0.76 | .937 |
| False rec | Word | 0.19 | 0.66 | 0.29 | .937 |
| | Pseudoword | 0.11 | 0.82 | 0.13 | .937 |
| | RFG | | | | |
| True rec | Word | -0.45 | 0.76 | -0.59 | .937 |
| | Pseudoword | 0.06 | 0.79 | 0.08 | .937 |
| False rec | Word | 0.15 | 0.74 | 0.21 | .937 |
| | Pseudoword | -0.51 | 0.72 | -0.70 | .937 |
| | LFG | | | | |
| True rec | Word | -0.94 | 0.67 | -1.40 | .486 |
| | Pseudoword | 1.04 | 1.02 | 1.02 | .596 |
| False rec | Word | 0.25 | 0.65 | 0.39 | .869 |
| | Pseudoword | -0.87 | 0.93 | -0.94 | .596 |
| | LvATL | | | | |

| | | | | | |
|---|---|---|---|---|---|
| True rec | Word | -0.02 | 1.12 | -0.02 | .984 |
| | Pseudoword | -0.20 | 0.78 | -0.26 | .869 |
| False rec | Word | 3.18 | 1.13 | 2.83 | .028 |
| | Pseudoword | 0.43 | 0.76 | 0.56 | .860 |
| | LvPFC | | | | |
| True rec | Word | -1.50 | 0.59 | -2.56 | .042 |
| | Pseudoword | -0.25 | 0.79 | -0.32 | .869 |
| False rec | Word | 1.68 | 0.55 | 3.07 | .025 |
| | Pseudoword | -0.84 | 0.70 | -1.20 | .553 |

**Table S3**. Contrasts between the estimated marginal means reflecting significant differences between Wfonts and Pfonts conditions for true and false visual recognition

| True recognition | ROI | ß | SE | z-value | p |
|---|---|---|---|---|---|
| | **Word - Pseudoword** | | | | |
| | **Visual model** | | | | |
| | EVC | -0.31 | 0.45 | -0.68 | .496 |
| | LG | -0.29 | 0.62 | -0.47 | .636 |
| | RFG | 1.28 | 1.11 | 1.16 | .496 |
| | LFG | -1.74 | 1.05 | -1.65 | .395 |
| | LvATL | 0.88 | 1.31 | 0.67 | .975 |
| | LvPFC | -0.21 | 0.72 | -0.30 | .809 |
| | **Visuosemantic model** | | | | |
| | EVC | 0.87 | 1.14 | 0.76 | .485 |
| | LG | 1.04 | 0.94 | 1.11 | .519 |
| | RFG | -0.81 | 0.99 | -0.81 | .424 |
| | LFG | -1.32 | 1.04 | -1.27 | .301 |
| | LvATL | -0.68 | 0.92 | -0.74 | .521 |
| | LvPFC | 0.45 | 1.00 | 0.45 | .838 |
| | **Semantic model** | | | | |
| | EVC | -1.06 | 1.22 | -0.87 | .769 |
| | LG | -1.35 | 1.09 | -1.24 | .693 |
| | RFG | -0.51 | 1.10 | -0.46 | .643 |
| | LFG | -1.98 | 1.22 | -1.62 | .269 |
| | LvATL | 0.18 | 1.36 | 0.13 | .895 |
| | LvPFC | -1.25 | 0.98 | -1.27 | .271 |
| False recognition | ROI | ß | SE | z-value | p |
| | **Word - Pseudoword** | | | | |
| | **Visual model** | | | | |
| | EVC | 0.45 | 0.49 | 0.92 | .478 |
| | LG | 0.74 | 0.66 | 1.11 | .356 |
| | RFG | -0.63 | 1.08 | -0.58 | .750 |
| | LFG | -0.10 | 1.00 | -0.10 | .924 |
| | LvATL | 0.94 | 1.31 | 0.71 | .975 |
| | LvPFC | -0.16 | 0.68 | -0.24 | .809 |
| | **Visuosemantic model** | | | | |
| | EVC | 1.44 | 1.01 | 1.43 | .451 |
| | LG | 0.77 | 0.89 | 0.86 | .519 |
| | RFG | 0.82 | 1.03 | 0.80 | .424 |
| | LFG | 1.18 | 0.97 | 1.21 | .301 |

| | | | | | |
|---|---|---|---|---|---|
| | LvATL | 1.30 | 0.87 | 1.49 | .291 |
| | LvPFC | 1.72 | 0.96 | 1.79 | .291 |
| | **Semantic model** | | | | |
| | EVC | 0.53 | 1.13 | 0.47 | .855 |
| | LG | 0.08 | 1.05 | 0.08 | .937 |
| | RFG | 0.66 | 1.03 | 0.64 | .643 |
| | LFG | 1.13 | 1.13 | 0.99 | .321 |
| | LvATL | 2.76 | 1.36 | 2.03 | .086 |
| | LvPFC | 2.52 | 0.89 | 2.84 | .009 |

**Table S4**. Contrasts between the estimated marginal means reflecting significant differences between true and false visual recognition for Wfonts and Pfonts conditions.

| Word | ROI | ß | SE | z-value | p |
|---|---|---|---|---|---|
| | **True rec – False rec** | | | | |
| | **Visual model** | | | | |
| | EVC | 1.80 | 0.45 | 3.98 | <.001 |
| | LG | 2.28 | 0.60 | 3.81 | <.001 |
| | RFG | 1.97 | 1.32 | 1.49 | .496 |
| | LFG | -0.46 | 1.02 | -0.45 | .867 |
| | LvATL | 0.05 | 1.56 | 0.03 | .975 |
| | LvPFC | -0.40 | 0.81 | -0.50 | .809 |
| | **Visuosemantic model** | | | | |
| | EVC | 0.74 | 1.07 | 0.70 | .485 |
| | LG | 0.70 | 0.78 | 0.89 | .519 |
| | RFG | 3.60 | 0.96 | 3.75 | <.001 |
| | LFG | -2.95 | 0.63 | -4.66 | <.001 |
| | LvATL | -0.47 | 0.73 | -0.64 | .521 |
| | LvPFC | -0.19 | 0.91 | -0.20 | .838 |
| | **Semantic model** | | | | |
| | EVC | -0.06 | 0.93 | -0.06 | .952 |
| | LG | -0.89 | 0.95 | -0.94 | .693 |
| | RFG | -0.60 | 1.06 | -0.57 | .643 |
| | LFG | -1.19 | 0.93 | -1.28 | .269 |
| | LvATL | -3.21 | 1.59 | -2.02 | .086 |
| | LvPFC | -3.19 | 0.80 | -3.97 | <.001 |
| Pseudoword | ROI | ß | SE | z-value | p |
| | **True rec – False rec** | | | | |
| | **Visual model** | | | | |
| | EVC | 2.56 | 0.49 | 5.22 | <.001 |
| | LG | 3.30 | 0.68 | 4.84 | <.001 |
| | RFG | 0.06 | 0.81 | 0.08 | .940 |
| | LFG | 1.18 | 1.03 | 1.15 | .504 |
| | LvATL | 0.10 | 1.00 | 0.10 | .975 |
| | LvPFC | -0.36 | 0.57 | -0.62 | .809 |
| | **Visuosemantic model** | | | | |
| | EVC | 1.32 | 1.09 | 1.21 | .451 |
| | LG | 0.43 | 1.03 | 0.42 | .677 |
| | RFG | 5.23 | 1.06 | 4.95 | .<001 |
| | LFG | -0.45 | 1.27 | -0.36 | .722 |

| | | | | | |
|---|---|---|---|---|---|
| | LvATL | 1.51 | 1.04 | 1.46 | .291 |
| | LvPFC | 1.09 | 1.04 | 1.05 | .590 |
| | **Semantic model** | | | | |
| | EVC | 1.53 | 1.38 | 1.11 | .769 |
| | LG | 0.54 | 1.18 | 0.46 | .863 |
| | RFG | 0.57 | 1.07 | 0.53 | .643 |
| | LFG | 1.91 | 1.38 | 1.38 | .269 |
| | LvATL | -0.63 | 1.09 | -0.58 | .752 |
| | LvPFC | 0.59 | 0.05 | 0.56 | .578 |

*ERS: Effects of reinstatement on visual true and false memory*

**Table S5.** Interaction of stimulus type and memory outcome on encoding-retrieval similarity (ERS). Paired t-tests comparing memory-related ERS differences (Hit > Miss; CR > FA) between Wfonts and Pfonts trials. Positive values reflect stronger ERS effects for the Wfonts condition; negative values indicate stronger effects for Pfonts. Cohen's *d* reports effect size.

| **True recognition** | | | | | | | |
|---|---|---|---|---|---|---|---|
| ROI | Condition | t(29) | p | d | Contrast | p | d |
| EVC | Word | 3.66 | .002 | 0.95 | -5.69 | <.001 | -1.24 |
| | Pseudoword | 12.86 | < .001 | 3.00 | | | |
| LG | Word | 3.62 | .002 | 1.04 | -3.51 | .013 | -0.93 |
| | Pseudoword | 14.74 | <.001 | 3.50 | | | |
| RFG | Word | 6.11 | <.001 | 1.78 | -7.03 | <.001 | -1.76 |
| | Pseudoword | 15.52 | <.001 | 3.93 | | | |
| LFG | Word | -0.43 | .873 | -0.13 | -0.62 | .796 | -0.17 |
| | Pseudoword | 0.49 | .700 | 0.13 | | | |
| LvATL | Word | 0.01 | .992 | 0.00 | -0.44 | .796 | -0.11 |
| | Pseudoword | 0.64 | .700 | 0.15 | | | |
| LvPFC | Word | 0.35 | .873 | 0.08 | 0.07 | .943 | 0.02 |
| | Pseudoword | 0.39 | .700 | 0.11 | | | |
| **False recognition** | | | | | | | |
| ROI | Condition | t(29) | p | d | Contrast | p | d |
| EVC | Word | 2.38 | .024 | 0.62 | 3.18 | .005 | 0.85 |
| | Pseudoword | 6.08 | <.001 | 1.58 | | | |
| LG | Word | 7.17 | <.001 | 1.69 | 4.88 | <.001 | 1.29 |
| | Pseudoword | 14.48 | <.001 | 4.15 | | | |
| RFG | Word | 12.54 | <.001 | 3.39 | -0.36 | .722 | -0.09 |
| | Pseudoword | 6.11 | <.001 | 1.66 | | | |
| LFG | Word | -9.38 | <.001 | -2.46 | 7.56 | <.001 | 2.29 |
| | Pseudoword | 3.51 | .002 | 0.91 | | | |
| LvATL | Word | -12.79 | <.001 | -3.93 | 6.03 | <.001 | 1.63 |
| | Pseudoword | -5.01 | <.001 | -1.18 | | | |
| LvPFC | Word | -5.73 | <.001 | -1.54 | 2.46 | .024 | 0.63 |

| | | | | |
|---|---|---|---|---|
| | Pseudoword | -0.91 | .370 | -0.24 |

*Mediation analysis: Hippocampal contributions to memory via pattern similarity*

**Table S6.** This table reports the average causal mediation effect (ACME) of hippocampal activation (posterior, anterior) on true recognition performance via encoding–retrieval similarity (ERS). Results are separated by ROI and stimulus condition (Word vs. Pseudoword). Confidence intervals (CI), standard errors (SE), uncorrected *p*-values, and FDR-corrected *p*-values are shown for each mediation pathway.

| **True recognition** | | | | | | | |
|---|---|---|---|---|---|---|---|
| ROI | Condition | Hippocampus | ACME | CI- | CI+ | p | FDR-p |
| EVC | Word | Posterior | -0.01 | -0.09 | 0.07 | .842 | .962 |
| EVC | Pseudoword | Posterior | 0.21 | 0.07 | 0.35 | .002 | .006 |
| LG | Word | Posterior | -0.02 | -0.07 | 0.04 | .588 | .962 |
| LG | Pseudoword | Posterior | -0.05 | -0.15 | 0.06 | .400 | .800 |
| RFG | Word | Posterior | 0.00 | -0.01 | 0.01 | .962 | .962 |
| RFG | Pseudoword | Posterior | 0.27 | 0.12 | 0.43 | <.000 | <.000 |
| LFG | Word | Posterior | 0.02 | -0.06 | 0.10 | .640 | .962 |
| LFG | Pseudoword | Posterior | 0.01 | -0.06 | 0.07 | .930 | .970 |
| LvATL | Word | Posterior | 0.00 | 0.00 | 0.00 | .948 | .962 |
| LvATL | Pseudoword | Posterior | 0.00 | -0.01 | 0.02 | .970 | .970 |
| LvPFC | Word | Posterior | -0.01 | -0.11 | 0.09 | .834 | .962 |
| LvPFC | Pseudoword | Posterior | 0.00 | -0.03 | 0.03 | .928 | .970 |
| EVC | Word | Anterior | 0.01 | -0.03 | 0.05 | .682 | .916 |
| EVC | Pseudoword | Anterior | 0.00 | -0.03 | 0.02 | .924 | .986 |
| LG | Word | Anterior | 0.00 | -0.05 | 0.04 | .902 | .916 |
| LG | Pseudoword | Anterior | 0.00 | -0.02 | 0.01 | .946 | .986 |
| RFG | Word | Anterior | 0.04 | -0.04 | 0.13 | .302 | .916 |
| RFG | Pseudoword | Anterior | -0.03 | -0.13 | 0.07 | .586 | .986 |
| LFG | Word | Anterior | 0.00 | -0.03 | 0.03 | .894 | .916 |
| LFG | Pseudoword | Anterior | 0.05 | -0.08 | 0.19 | .422 | .986 |
| LvATL | Word | Anterior | 0.00 | -0.02 | 0.02 | .916 | .916 |
| LvATL | Pseudoword | Anterior | 0.00 | -0.05 | 0.05 | .986 | .986 |
| LvPFC | Word | Anterior | -0.01 | -0.05 | 0.03 | .790 | .916 |
| LvPFC | Pseudoword | Anterior | -0.01 | -0.06 | 0.04 | .860 | .986 |

**Table S7.** This table reports the average causal mediation effect (ACME) of hippocampal activation (posterior, anterior) on false recognition performance via encoding–retrieval similarity (ERS). Results are separated by ROI and stimulus condition (Word vs. Pseudoword). Confidence intervals (CI), standard errors (SE), uncorrected *p*-values, and FDR-corrected *p*-values are shown for each mediation pathway.

| **False recognition** |
|---|

| ROI | Condition | Hippocampus | ACME | CI- | CI+ | p | FDR-p |
|---|---|---|---|---|---|---|---|
| EVC | Word | Posterior | -0.16 | -0.28 | -0.03 | .004 | .012 |
| EVC | Pseudoword | Posterior | 0.04 | -0.05 | 0.13 | .434 | .713 |
| LG | Word | Posterior | 0.01 | -0.04 | 0.07 | .788 | .890 |
| LG | Pseudoword | Posterior | 0.02 | -0.05 | 0.10 | .594 | .713 |
| RFG | Word | Posterior | -0.23 | -0.35 | -0.11 | <.000 | <.000 |
| RFG | Pseudoword | Posterior | 0.03 | -0.05 | 0.10 | .534 | .713 |
| LFG | Word | Posterior | -0.06 | -0.20 | 0.07 | .340 | .510 |
| LFG | Pseudoword | Posterior | -0.04 | -0.11 | 0.04 | .402 | .713 |
| LvATL | Word | Posterior | 0.00 | -0.03 | 0.04 | .890 | .890 |
| LvATL | Pseudoword | Posterior | 0.00 | -0.08 | 0.09 | .996 | .996 |
| LvPFC | Word | Posterior | -0.07 | -0.20 | 0.06 | .296 | .510 |
| LvPFC | Pseudoword | Posterior | -0.04 | -0.14 | 0.05 | .340 | .713 |
| EVC | Word | Anterior | 0.06 | -0.06 | 0.18 | .344 | .958 |
| EVC | Pseudoword | Anterior | 0.01 | -0.07 | 0.09 | .808 | .980 |
| LG | Word | Anterior | -0.04 | -0.20 | 0.11 | .538 | .958 |
| LG | Pseudoword | Anterior | 0.00 | -0.09 | 0.09 | .980 | .980 |
| RFG | Word | Anterior | 0.02 | -0.05 | 0.08 | .680 | .958 |
| RFG | Pseudoword | Anterior | 0.00 | -0.01 | 0.01 | .960 | .980 |
| LFG | Word | Anterior | 0.01 | -0.03 | 0.04 | .958 | .958 |
| LFG | Pseudoword | Anterior | -0.20 | -0.35 | -0.04 | .010 | .030 |
| LvATL | Word | Anterior | -0.07 | -0.21 | 0.06 | .250 | .958 |
| LvATL | Pseudoword | Anterior | -0.18 | -0.34 | -0.02 | .010 | .030 |
| LvPFC | Word | Anterior | 0.01 | -0.04 | 0.05 | .896 | .958 |
| LvPFC | Pseudoword | Anterior | 0.00 | -0.04 | 0.05 | .856 | .980 |

### Materials and Methods

#### *Image preprocessing*

Data processing was performed using fMRIPrep 23.0.1 (5), which is based on Nipype 1.8.5 (6). Fieldmaps were collected for each subject. A B0-nonuniformity map (or fieldmap) was estimated based on two (or more) echo-planar imaging (EPI) references with topup (7). Then, all T1-weighted (T1w) images were corrected for intensity non-uniformity (INU) with N4BiasFieldCorrection (8), distributed with ANTs 2.3.3 (9). The T1w-reference was then skull-stripped with a Nipype implementation of the antsBrainExtraction.sh workflow (from ANTs), using OASIS30ANTs as target template. Brain tissue segmentation of cerebrospinal fluid (CSF), white-matter (WM) and gray-matter (GM) was performed on the brain-extracted T1w using fast (FSL 6.0.5.1:57b01774; (10)). An anatomical T1w-reference map was computed after registration of 2 T1w images (after INU-correction) using mri_robust_template (FreeSurfer 7.3.2; (11)). Volume-based spatial normalization to one standard space (MNI152NLin2009cAsym) was performed through nonlinear registration with antsRegistration (ANTs 2.3.3), using brain-extracted versions of both T1w reference and the T1w template. The following template was selected for spatial normalization and accessed with TemplateFlow (23.0.0; (12): ICBM 152 Nonlinear Asymmetrical template version 2009c (13). For each of the 14 BOLD runs found per subject (across sessions), the following preprocessing was performed. First, a reference volume and its skull-stripped version were generated using a custom methodology of fMRIPrep. Head-motion parameters with respect to the BOLD reference (transformation matrices, and six corresponding rotation and translation parameters) are estimated before any spatiotemporal filtering using mcflirt (FSL 6.0.5.1:57b01774; (14). The estimated fieldmap was then aligned with rigid-registration to the target EPI (echo-planar imaging) reference run. The field coefficients were mapped on to the reference EPI using the transform. BOLD runs were slice-time corrected to 0.346s (0.5 of slice acquisition range 0s-0.693s) using 3dTshift from AFNI (15). The BOLD reference was then co-registered to the T1w reference using mri_coreg (FreeSurfer) followed by flirt (FSL 6.0.5.1:57b01774; (16)) with the boundary-based registration (17) cost-function. Co-registration was configured with six degrees of freedom. The BOLD time-series were resampled into standard space, generating a preprocessed BOLD run in MNI152NLin2009cAsym space. First, a reference volume and its skull-stripped version were generated using a custom methodology of fMRIPrep. All resamplings can be performed with a single interpolation step by composing all the pertinent transformations (i.e. head-motion transform matrices, susceptibility distortion correction when available, and co-registrations to anatomical and output spaces). Gridded (volumetric) resamplings were performed using antsApplyTransforms (ANTs), configured with Lanczos interpolation to minimize the smoothing effects of other kernels (18). Non-gridded (surface) resamplings were performed using mri_vol2surf (FreeSurfer). For RSA, we used unsmoothed preprocessed BOLD time series in original space to keep the finer-grained structure of activity.

*ROI*

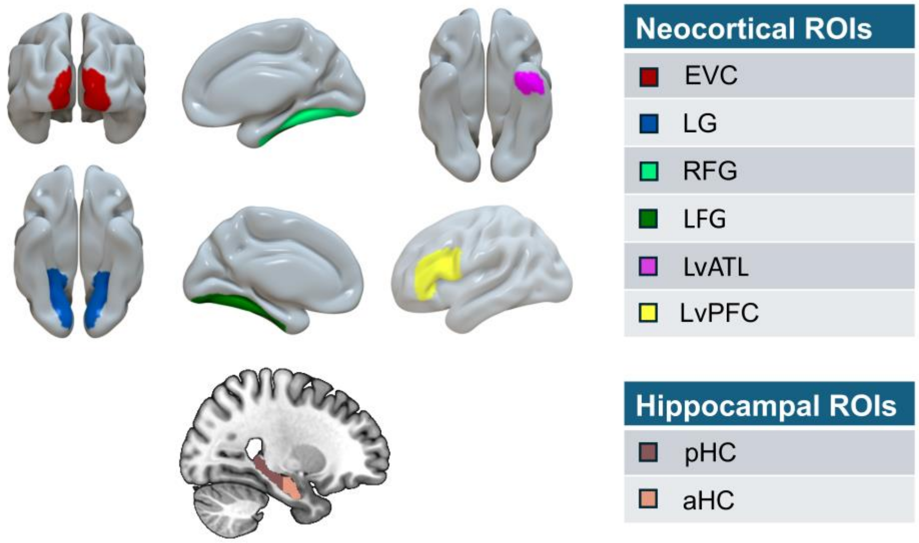


**Figure S3.** Regions of interest (ROIs). Neocortical ROIs included early visual cortex (EVC; BA17/18), lingual gyrus (LG), right fusiform gyrus (rFG), left fusiform gyrus (lFG), left ventral anterior temporal lobe (lvATL), and left inferior frontal gyrus (LIFG; BA44/45). Hippocampal ROIs were subdivided along the anterior–posterior axis into posterior hippocampus (pHPC) and anterior hippocampus (aHPC). ROIs were defined in standard MNI space and transformed to individual functional space. Visualization was performed using Surf Ice and MRIcroGL.
